# Rho signaling promotes cell excitability and synaptic transmission

**DOI:** 10.1101/2025.11.13.688363

**Authors:** Ariana José, Pravat Dhakal, Rossana Signorelli, Caroline Case, Sana I. Chaudhry, Kevin M. Collins

## Abstract

Gα_q_ signaling through Trio RhoGEF and Phospholipase C effectors promotes neurotransmitter release at synapses. Whether activated Rho and Phospholipase C signaling acts through shared or independent pathways remains unclear. We use the egg-laying circuit of *C. elegans*, which is regulated positively by Gα_q_ signaling and negatively by Gα_o_ signaling, to determine the role of Rho GTPase in G-protein regulation of neural circuit excitability. We previously showed that Trio signals in both the presynaptic Hermaphrodite-Specific command Neurons (HSNs) and the postsynaptic egg-laying vulval muscles they innervate to promote egg laying. Here we show that its effector Rho similarly signals both pre- and post-synaptically. Expression of constitutively active Rho-1(G14V) in either the HSNs or vulval muscles increased their Ca^2+^ activity and stimulated egg laying. Conversely, reducing Rho signaling in the HSNs via expression of Rho-1(T19N) or C3 transferase expression inhibited HSN Ca^2+^ activity and egg laying. Animals lacking Rho function in HSN still had vulval muscle Ca^2+^ activity that supported contraction and egg release, although this Ca^2+^ activity lacked the enhancement associated with HSN neurotransmitter release, suggesting it was largely driven by stretch-dependent feedback. Together, these results show that Rho signals in both the HSNs and vulval muscles to promote excitability and egg laying.

**Article Summary:** Rho-type G-proteins have been extensively studied for their role in neural development, but accumulating evidence points to a role in regulating synaptic transmission and neural circuit activity. Using powerful *in vivo* imaging and genetic techniques in the nematode *Caenorhabditis elegans*, the authors show that Rho signals in both presynaptic neurons and postsynaptic muscles to promote circuit activity and behavior. These findings solidify Rho as a key regulator of neurotransmitter signaling.

## Introduction

Rho GTPases, a family of small G proteins in the Ras superfamily, serve as molecular switches to transduce signaling from extracellular ligands to intracellular effectors (Nobes and Hall, 1995; Ridley and Hall, 1992; Ridley et al., 1992). Studies have shown that Rho GTPases regulate cytoskeletal reorganization (Nobes and Hall, 1995; Ridley and Hall, 1992). They also play a pivotal role in several other aspects of cell biology including membrane transport pathways, and transcription factor activation, which has a significant role in the development and function of the nervous system (Gallo and Letourneau, 1998; Govek et al., 2005). Rho GTPases are activated by a large class of guanine nucleotide exchange factors (GEFs) of which Trio is a prominent member. Mutations in the human ortholog of Trio cause developmental defects including autism (Eid et al., 2025; Tian et al., 2021), and experiments reducing Trio and effector functions in mice, *Aplysia*, flies, and worms lead to a variety of developmental or behavioral defects (Bateman et al., 2000; Carrizales et al., 2025; Ishchenko et al., 2025; Steven et al., 1998; Williams et al., 2007).

Trio contains two DH/PH GEF domains (Debant et al., 1996; Steven et al., 1998). The first GEF domain activates Rac whose signaling in neurons contributes to axon guidance (Bateman et al., 2000; Steven et al., 1998). Recent work shows that Rac1 also functions in neurons to reduce synaptic transmission, possibly through altered actin reorganization at synapses (Keine et al., 2022). The second Trio GEF domain activates Rho, and is a direct effector for activation by the heterotrimeric GTPase Gα_q_ (Rojas et al., 2007; Williams et al., 2007). Trio and other RhoGEF effectors are present at synapses and regulate synaptic transmission (Doussau et al., 2000; Herring and Nicoll, 2016; Terry-Lorenzo et al., 2016). Rho is found on rat brain synaptosome fractions, and toxins that block Rho-family proteins including Rac reduce acetylcholine release from neurons and paired-pulse facilitation in *Aplysia* (Doussau et al., 2000). The Rho activator Trio promotes AMPA receptor synaptic transmission and long-term potentiation, and this may be enhanced by CaMKII phosphorylation (Herring and Nicoll, 2016). Work in *C. elegans* has similarly shown that Gα_q_, Trio, and Rho-1 signaling promote neurotransmission (Brundage et al., 1996; Hiley et al., 2006; McMullan et al., 2006; Williams et al., 2007), and determining how and where Rho-1 functions would inform our understanding of how Gα_q_ signaling through Trio and Rho modulates synaptic plasticity.

Another major effector for Gα_q_ is PLCß which generates the second messengers IP_3_ and DAG to promote synaptic transmission (Lackner et al., 1999; Miller et al., 1999; Smrcka et al., 1991; Taylor et al., 1991). Gα_q_ signaling through PLCß in the *Drosophila* visual system leads to DAG production, activation of TRP channels, and depolarization of the photoreceptors (Bloomquist et al., 1988; Niemeyer et al., 1996; Sturgeon and Magoski, 2018). In *C. elegans*, DAG-mimetic phorbol esters promote cell Ca^2+^ activity and rescue behavior defects of Trio RhoGEF and PLCß mutants (Dhakal et al., 2022; Nurrish et al., 1999; Williams et al., 2007), suggesting DAG-regulated effectors and/or ion channels are downstream of both Rho and PLCß. Rho can activate PLCε which may explain how signaling through separate Trio and PLCß effectors may ultimately converge to generate IP_3_ and DAG to regulate cell excitability (Kariya et al., 2004; Seifert et al., 2008). NALCN channels have been identified as putative targets for modulation by G protein signaling in the mammalian brain (Lutas et al., 2016) and act downstream of Gα_q_, Trio, and Rho in the control of worm locomotion (Topalidou et al., 2017a; Topalidou et al., 2017b). NALCN channels are also expressed in the *C. elegans* egg-laying circuit, and mutations that increase NALCN channel activity stimulate egg laying (Yeh et al., 2008). However, animals lacking NALCN channels have grossly normal egg-laying behavior and pharmacological responses to serotonin and phorbol esters (Jose and Collins, 2024), suggesting NALCN channels are not exclusive targets for serotonin signaling through Gα_q_ and Rho signaling through DAG in *C. elegans*.

Rho-1 is an essential gene in *C. elegans*, and methods to manipulate Rho-1 signaling cell-specifically have shown how it regulates synaptic transmission and behavior. Expression of the GTP-locked, constitutively active Rho-1(G14V) mutant increases locomotion and egg-laying behavior, while expression of C3 transferase, which ribosylates ADP and specifically inhibits Rho-1 function, reduced these behaviors (Hiley et al., 2006; McMullan and Nurrish, 2011; McMullan et al., 2006). Activated Rho-1 binds to and inhibits Diacylglycerol (DAG) kinase (McMullan et al., 2006), blocking conversion of membrane DAG to phosphatidic acid (Jose and Koelle, 2005). Genetic loss of DAG Kinase in *C. elegans* is thought to increase synaptic DAG levels, potentiating neurotransmitter release (Jose and Koelle, 2005; Nurrish et al., 1999). DAG recruits and activates downstream molecules including UNC-13 which promote acetylcholine neurotransmitter release at the neuromuscular junction (Lackner et al., 1999). Gα_q_ signals in both the presynaptic Hermaphrodite Specific Neurons (HSNs) and the postsynaptic vulval muscles they innervate to regulate serotonin transmission and egg-laying behavior (Bastiani et al., 2003; Dhakal et al., 2022; Tanis et al., 2008). Loss of Gα_q_ and Trio RhoGEF signaling blocks egg laying (Brundage et al., 1996; Williams et al., 2007), leading to deficiencies in cell excitability that resemble the consequences of too much inhibitory Gα_o_ signaling (Koelle and Horvitz, 1996; Kumar et al., 2021; Ravi et al., 2021). Gα_o_ signaling opposes excitatory Gα_q_ signaling and depresses the electrical excitability and Ca^2+^ activity of the HSN command neurons (Ravi et al., 2021), allowing for sensory and stretch-dependent regulation of egg laying (Medrano et al., 2025; Ringstad and Horvitz, 2008). Determining whether changes in Rho-1 signaling cause similar changes in HSN and vulval muscle Ca^2+^ activity would help explain how Gα_o_ antagonizes Gα_q_ signaling to regulate cell excitability and synaptic transmission.

Here, we genetically manipulate Rho signaling in the *C. elegans* egg-laying circuit to study where and how Rho regulates neurotransmission and behavior. We find that Rho signals both presynaptically in the HSNs and postsynaptically in the vulval muscles to promote egg laying. Using ratiometric Ca^2+^ imaging in behaving animals, we find that Rho signaling promotes HSN excitability and synaptic transmission. This work suggests Gα_q_ signaling through Rho modulates ion channels to promote neuron and muscle electrical excitability.

## Materials and Methods

### Strains

*C. elegans* worms were maintained at 20°C on Nematode Growth Medium (NGM) agar plates with *E. coli* OP50 as a source of food as described previously (Brenner, 1974). All behavior assays and fluorescence imaging experiments were performed with age-matched adult hermaphrodites aged 24-36 h after the late L4 stage. All strains used in this study are listed in **Table 1**.

**Table 1:**
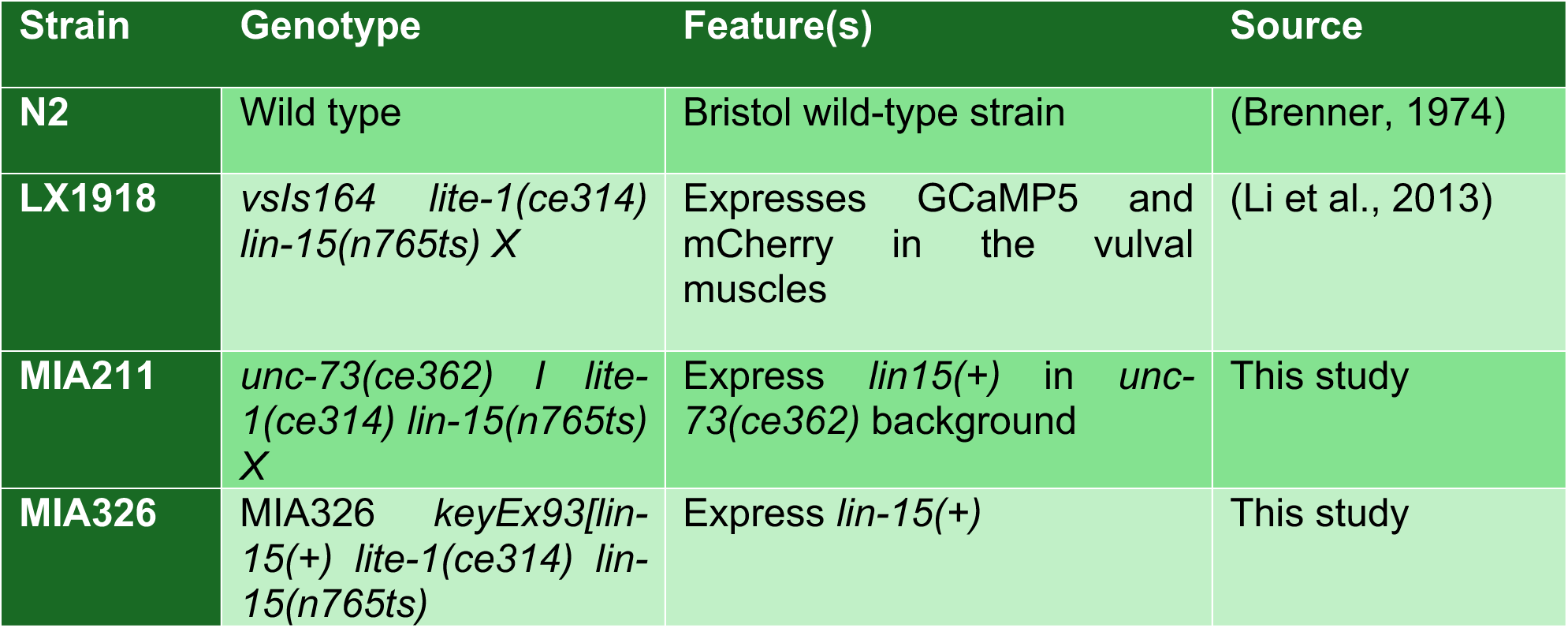
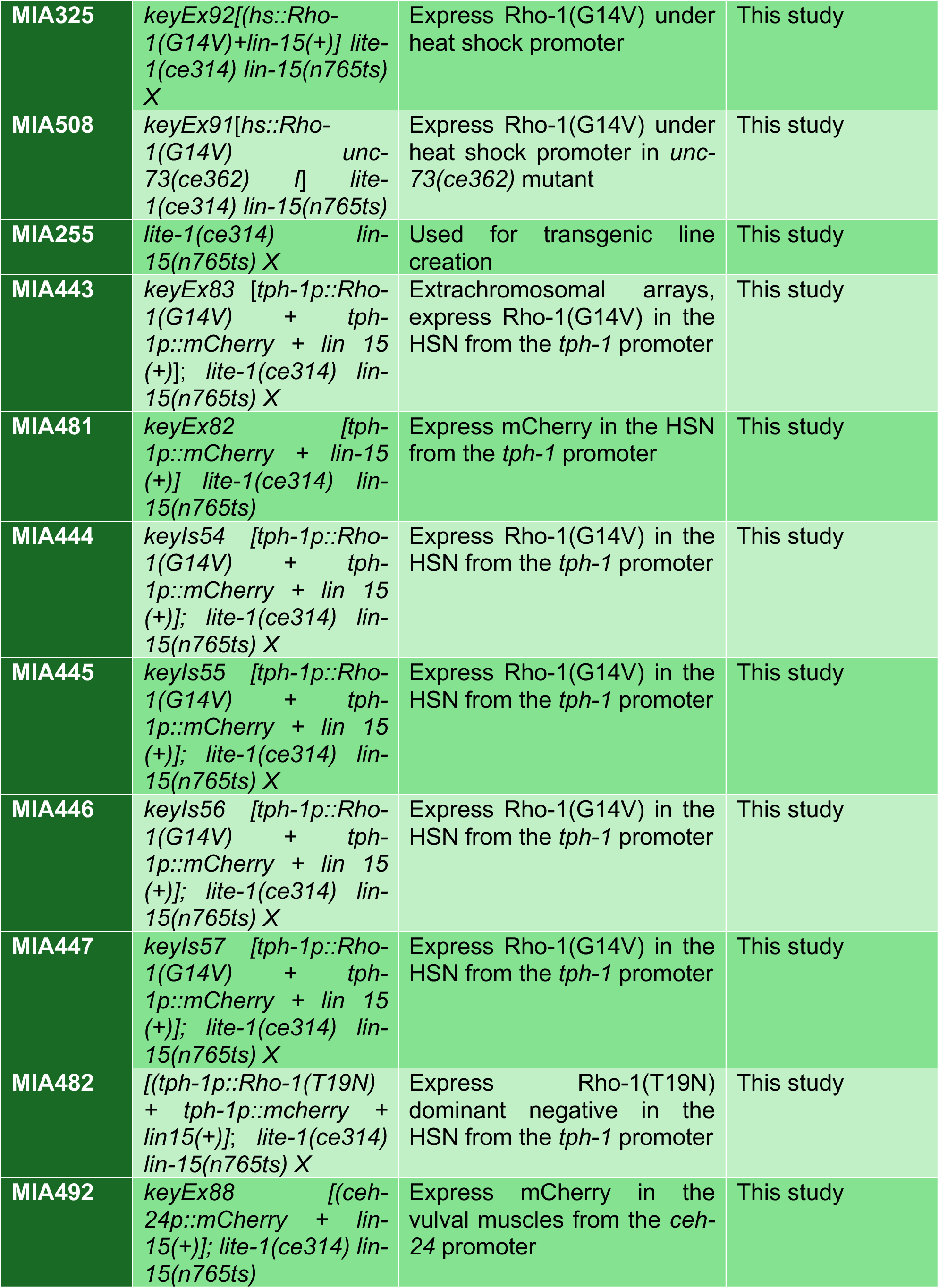
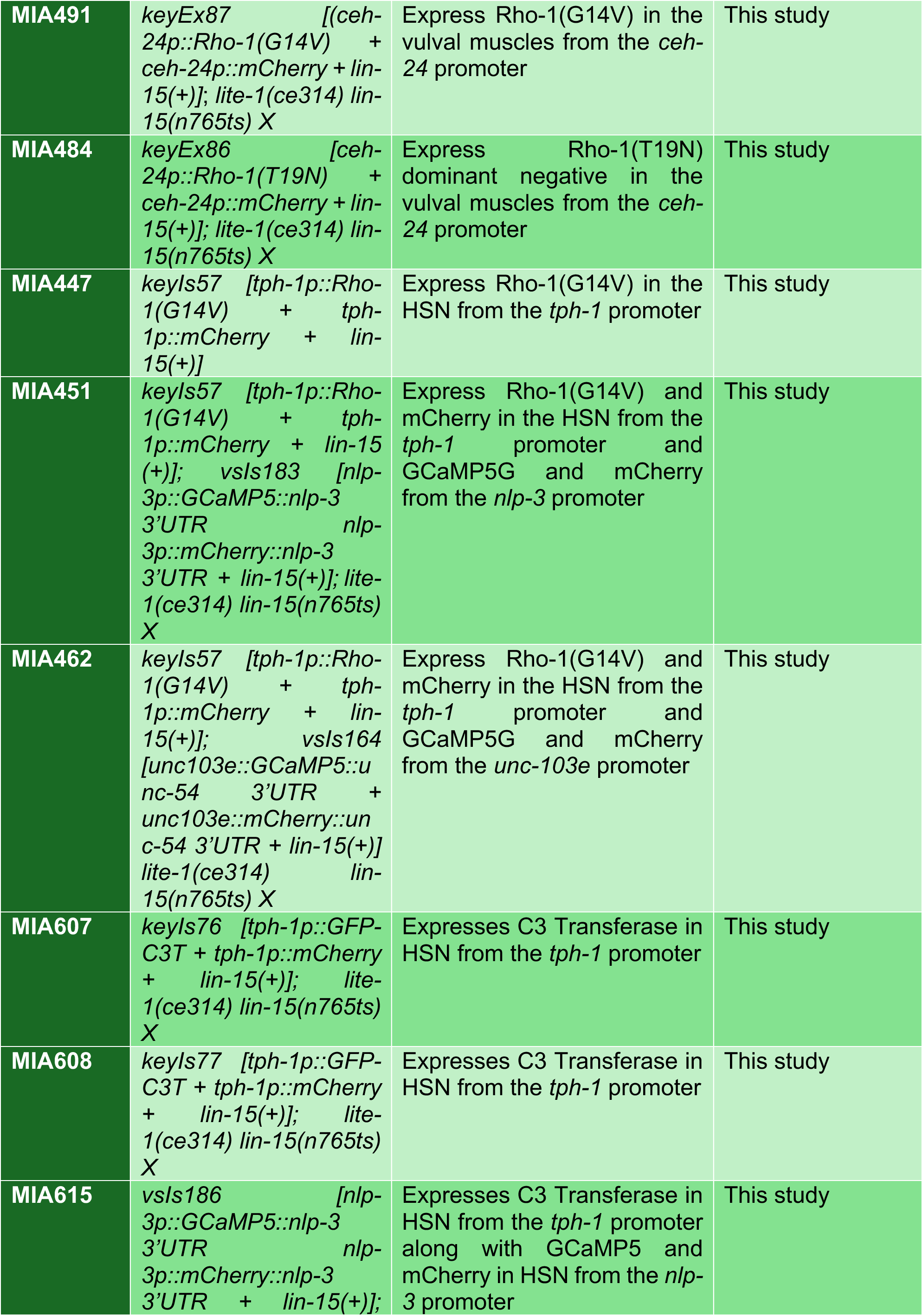
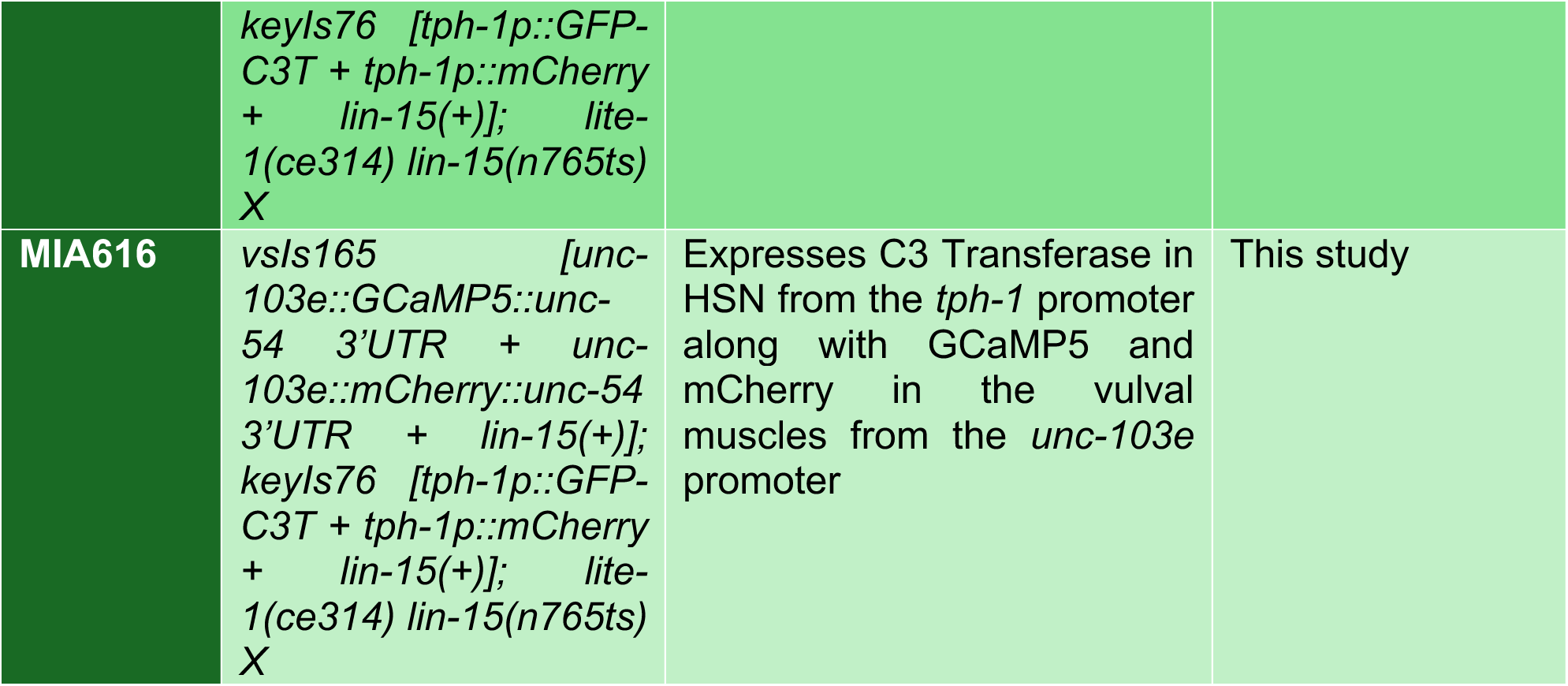
List of strains used in this study.

### Molecular biology and transgenes

#### Rho GTPase transgenes

Plasmid DNA was microinjected into gonads of adult hermaphrodites to generate transgenes. Plasmid pQT#42 containing sequences that express RHO-1(G14V) behind the heat-shock promoter was a gift of Joshua Kaplan and Stephen Nurrish, Massachusetts General Hospital, Boston, MA (McMullan et al., 2006). Plasmid pL15EK (50ng/µL) alone or along with pQT#42 (5ng/µL) was injected into LX1832 *lite-1(ce314) lin-15(n765ts) X* animals to generate extrachromosomal arrays. Five independent transgenic lines were used for behavior experiments from which a transgenic line (MIA326 *keyEx93[lin-15(+) lite-1(ce314) lin-15(n765ts)* and MIA325 *keyEx92 [(hs::Rho-1(G14V) + lin-15(+)] lite-1(ce314) lin-15(n765ts) X* respectively) was kept. To express RHO-1(G14V) in *unc-73(ce362)* plasmid pL15EK (50 ng/µL) along with pQT#42 (5 ng/µL) was injected into MIA211 *unc-73(ce362) I; lite-1(ce314) lin-15(n765ts) X* animals to generate extrachromosomal arrays. One independent transgenic line MIA508 *keyEx91* [*hs::Rho-1(G14V) lin-15(+)*] *unc-73(ce362) I; lite-1(ce314) lin-15(n765ts)* was used for behavior experiments.

To express RHO-1(G14V) in HSN, pSIC55 *[tph-1p::Rho-1(G14V)]* was generated by the FastCloning method, as described (Li et al., 2011). Briefly, PCR amplification of vector template from pKMC42 (*tph-1p::mCherry*) and insert template from pQT#42 (*hs::Rho-1(G14V))* was generated using oligos listed in **Table 2**. Vector and insert amplicons were mixed, digested with DpnI, transformed into DH5α bacteria, generating pSIC55. Plasmids pKMC42 *(tph-1p::mCherry,* 80 ng/µl) and pL15EK (50 ng/µL) alone or along with pSIC55 (80 ng/µL) was injected into MIA255 *lite-1(ce314) lin-15(n765ts) X* hermaphrodites to create five independent extrachromosomal arrays from which single transgenic lines (MIA443 *keyEx83*[*tph-1p::Rho-1(G14V) + tph-1p::mCherry + lin-15(+)*] *lite-1(ce314) lin-15(n765ts) X* and MIA481 *keyEx82 [tph-1p::mCherry + lin-15(+)] lite-1(ce314) lin-15(n765ts),* respectively, were kept*. keyEx83* was then integrated into chromosomes using UV/Trimethylpsoralen. Four independent integrants, *keyIs54-57 [tph-1p::Rho-1(G14V) + tph-1p::mCherry + lin 15 (+)*]*; lite-1(ce314) lin-15(n765ts) X.* were recovered and strains carrying them were backcrossed six times to MIA255 *lite-1(ce314) lin-15(n765ts) X,* generating lines MIA444-447, respectively. One integrated strain was subsequently crossed with other strains for HSN and vulval muscle Ca^2+^ imaging (see below). Plasmid pSIC55 [*tph-1p::(Rho-1(G14V*)] was also used as a template for QuikChange mutagenesis (Braman et al., 1996; Edelheit et al., 2009) to introduce the Rho-1(T19N) mutant generating pRS3*(tph-1p::Rho-1(T19N)* using oligonucleotides listed in **Table 2**. Plasmids pKMC42 *(tph-1p::mCherry,* 80 ng/µl) and pL15EK (50 ng/µL) alone or along with pRS3 (80ng/µL) was injected into MIA255 *lite-1(ce314) lin-15(n765ts) X* animals to create five independent extrachromosomal arrays from which a single transgenic line MIA482 *keyEx84 [(tph-1p::Rho-1(T19N) + tph-1p::mCherry + lin15(+); lite-1(ce314) lin-15(n765ts) X* was kept.

**Table 2:**
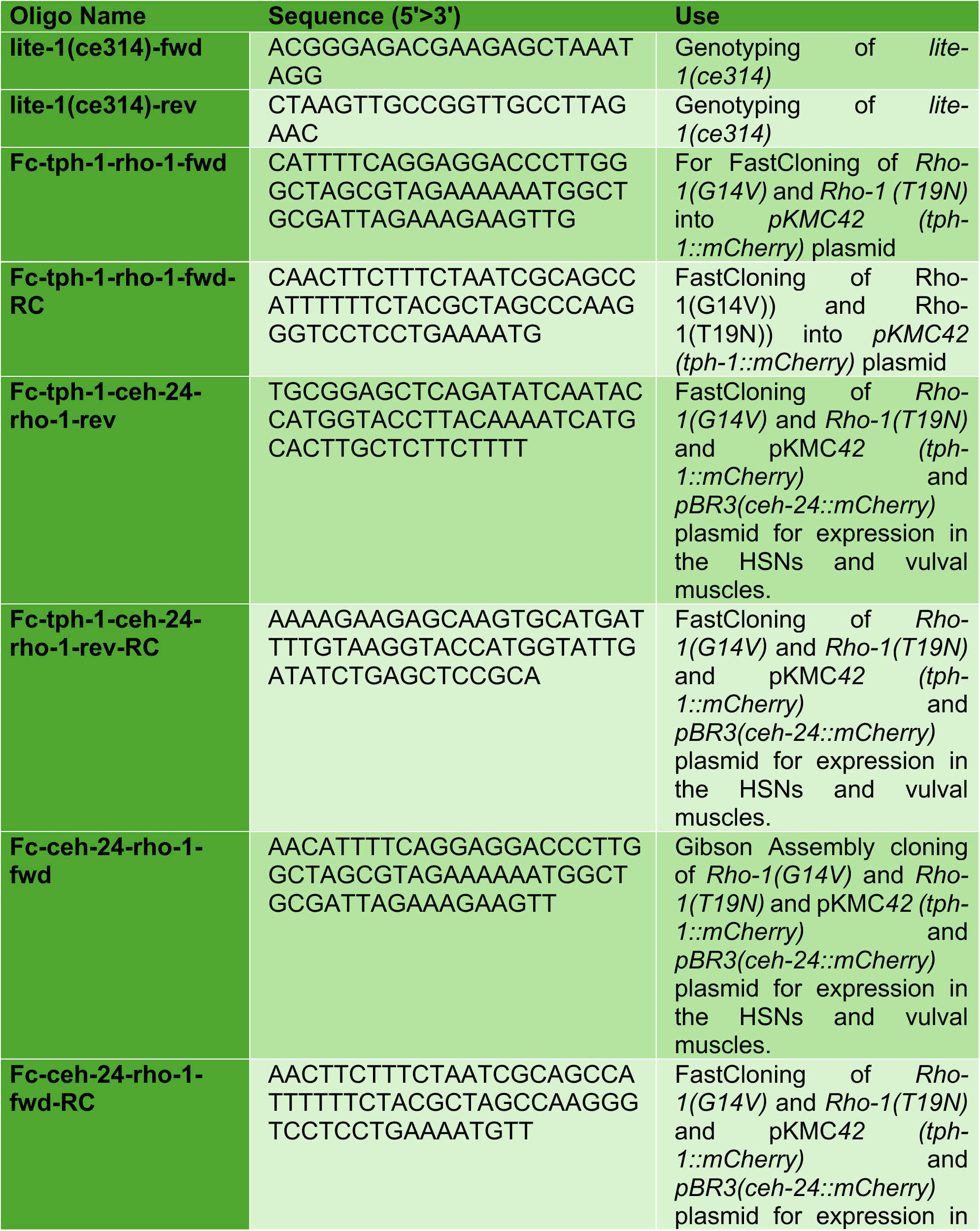
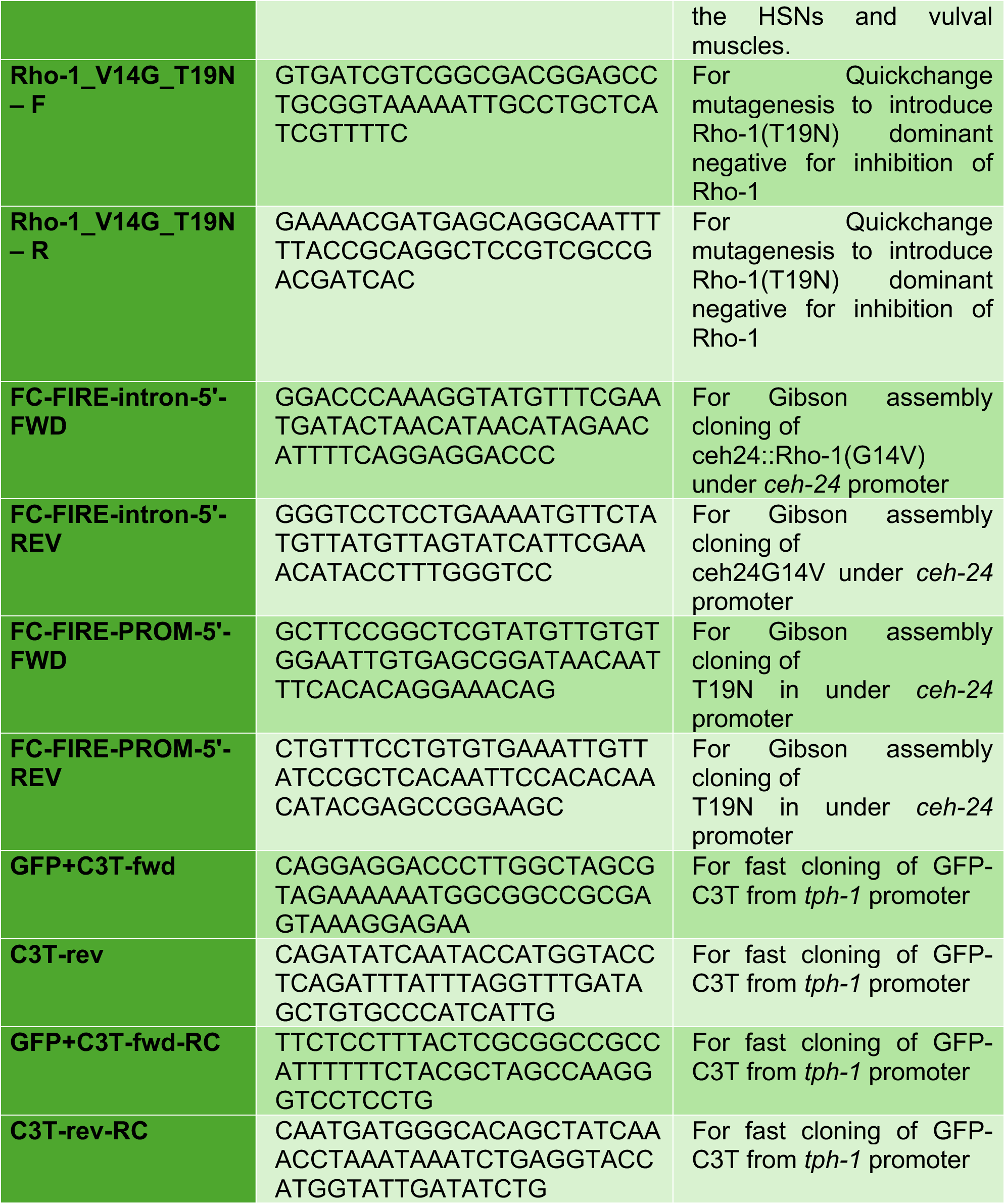
Lists of oligonucleotide sequences used in this study.

To express RHO-1(G14V) and RHO-1(T19N) in the vulval muscles from the *ceh-24* promoter, pRS4 (*ceh-24p::Rho-1(G14V)* and pRS5 *(ceh24p::Rho-1(T19N),* respectively were generated using Gibson assembly cloning from plasmid pSIC55 as a template. Plasmids pBR3 *(ceh-24p::mCherry,* 5ng/µl) and pL15EK (50 ng/µL) alone or along with pRS4 (5 ng/μl) or pRS5 (10 ng/µl) were injected into MIA255 animals to create five independent extrachromosomal arrays from which a single transgenic line from each (MIA484 *keyEx86 [ceh-24p::Rho-1(T19N) + ceh-24p::mCherry + lin-15(+)]; lite-1(ce314) lin-15(n765ts) X,* MIA491 *keyEx87 [(ceh-24p::Rho-1(G14V) + ceh-24p::mCherry + lin-15(+)*; *lite-1(ce314) lin-15(n765ts) X* and MIA492 *keyEx88 [(ceh-24p::mCherry + lin-15(+)]; lite-1(ce314) lin-15(n765ts)*, respectively) was kept.

To express GFP-fused C3 transferase (C3T) enzyme in the HSNs, pAEJ15 (*tph-1p::GFP:C3T)* was generated by FastCloning. Briefly, PCR amplification of vector template from pKMC42 (*tph-1p::mCherry*) and insert template from pQT#99 (*hs::GFP-C3T)* was generated using oligonucleotides listed in **Table 2**. Vector and insert amplicons were mixed, digested with DpnI, transformed into DH5α bacteria, generating pAEJ15. Plasmids pKMC42 *(tph-1p::mCherry,* 90 ng/µl) and pL15EK (50 ng/µL) alone or along with pAEJ15 (20 ng/µL) was injected into LX1832 *lite-1(ce314) lin-15(n765ts) X* hermaphrodites to create three independent extrachromosomal arrays from which single transgenic lines (MIA576 *keyEx98* [*tph-1p::GFP-C3T+ tph-1p::mCherry + lin-15(+)*] *lite-1(ce314) lin-15(n765ts) X* and MIA577 *keyEx99 [tph-1p::mCherry + lin-15(+)] lite-1(ce314) lin-15(n765ts),* respectively, were kept*. keyEx98* was then integrated into chromosomes using UV/Trimethylpsoralen. Two independent integrants, *keyIs76 and keyIs77 [tph-1p::GFP-C3T + tph-1p::mCherry + lin-15(+)*]*; lite-1(ce314) lin-15(n765ts) X,* were recovered, and strains carrying them were backcrossed six times to LX1832 *lite-1(ce314) lin-15(n765ts) X,* generating lines MIA607 and MIA608, respectively. Strain MIA607 was subsequently crossed with other strains for HSN and vulval muscle Ca^2+^ imaging.

HSN and vulval muscle Ca^2+^ activity was recorded using GCaMP5G (Akerboom et al., 2012) which was expressed along with mCherry from the *nlp-3* and *unc-103e* promoters, respectively, as previously described (Collins and Koelle, 2013; Collins et al., 2016; Ravi et al., 2018a). To visualize HSN Ca^2+^ activity in Rho-1(G14V) mutants, LX2004 *vsIs183 [nlp-3p::GCaMP5::nlp-3 3’UTR nlp-3p::mCherry::nlp-3 3’UTR + lin-15(+)] lite-1(ce314) lin-15(n765ts) X* males were crossed with MIA447 *keyIs57 [tph-1p::Rho-1(G14V) + tph-1p::mCherry + lin-15 (+)]; lite-1(ce314) lin-15(n765ts) X* hermaphrodites, and the fluorescent cross-progeny were picked and allowed to self. *keyIs57* and *vsIs183* were then homozygosed, generating MIA451 *keyIs57 [tph-1p::Rho-1(G14V) + tph-1p::mCherry + lin-15 (+)]; vsIs183 [nlp-3p::GCaMP5::nlp-3 3’UTR nlp-3p::mCherry::nlp-3 3’UTR + lin-15(+)]; lite-1(ce314) lin-15(n765ts) X* strains, respectively. The presence of *keyIs57* was confirmed by its hyperactive egg-laying phenotype; *vsIs183* was confirmed using mCherry fluorescence, and *lite-1(ce314)* was confirmed with PCR genotyping. For vulval muscle Ca^2+^ activity measurements, LX1918 *vsIs164 [unc-103e::GCaMP5::unc-54 3’UTR + unc-103e::mCherry::unc-54 3’UTR + lin-15(+)] lite-1(ce314) lin-15(n765ts) X* (Collins et al., 2016) males were crossed with MIA447 *keyIs57 [tph-1p::Rho-1(G14V) + tph-1p::mCherry + lin-15 (+)]; lite-1(ce314) lin-15(n765ts) X* hermaphrodites, and the fluorescent cross-progeny were picked and allowed to self. F2 hermaphrodites bearing *keyIs57* and *vsIs164* were homozygosed, generating MIA462 *keyIs57 [tph-1p::Rho-1(G14V) + tph-1p::mCherry + lin-15(+)]; vsIs164[unc-103e::GCaMP5::unc-54 3’UTR + unc-103e::mCherry::unc-54 3’UTR + lin-15(+)] lite-1(ce314) lin-15(n765ts) X.* Presence of *keyIs57* was confirmed by its hyperactive egg-laying phenotype, *vsIs164* was confirmed through mCherry expression, and *lite-1(ce314)* was confirmed with PCR genotyping.

To visualize HSN Ca^2+^ activity in GFP-C3T transgenic lines, LX2007 *vsIs186 [nlp-3p::GCaMP5::nlp-3 3’UTR nlp-3p::mCherry::nlp-3 3’UTR + lin-15(+)]; lite-1(ce314) lin-15(n765ts) X* (Collins et al., 2016) males were crossed with MIA607 *keyIs76 [tph-1p::GFP-C3T + tph-1p::mCherry + lin-15 (+)]; lite-1(ce314) lin-15(n765ts) X* hermaphrodites, and the fluorescent cross-progeny were picked and allowed to self. *keyIs76* and *vsIs186* were then homozygosed, generating the MIA616 *keyIs76 [tph-1p::GFP-C3T + tph-1p::mCherry + lin-15 (+)]; vsIs186 [nlp-3p::GCaMP5::nlp-3 3’UTR nlp-3p::mCherry::nlp-3 3’UTR + lin-15(+)]; lin-15(n765ts) X* strain. The presence of *keyIs76* was confirmed by its defective egg-laying phenotype; *vsIs186* was confirmed using mCherry fluorescence. For vulval muscle Ca^2+^ activity measurements, LX1919 *vsIs165 [unc-103e::GCaMP5::unc-54 3’UTR + unc-103e::mCherry::unc-54 3’UTR + lin-15(+)]; lite-1(ce314) lin-15(n765ts) X* (Collins et al., 2016) males were crossed with MIA607 *keyIs76 [tph-1p::GFP-C3T + tph-1p::mCherry + lin-15(+)]; lite-1(ce314) lin-15(n765ts) X* hermaphrodites, and the fluorescent cross-progeny were picked and allowed to self. F2 hermaphrodites bearing *keyIs76* and *vsIs165* were homogygosed, generating MIA615 *keyIs76 [tph-1p::GFP-C3T + tph-1p::mCherry + lin-15(+)]; vsIs165 [unc-103e::GCaMP5::unc-54 3’UTR + unc-103e::mCherry::unc-54 3’UTR + lin-15(+)] lin-15(n765ts) X.* Presence of *keyIs76* was confirmed by its defective egg-laying phenotype, and *vsIs165* was confirmed through mCherry expression.

### Behavior assays

Quantification of egg retention was performed as described (Chase and Koelle, 2004). Staged adults were obtained by picking late L4 animals, culturing them 24-30 h at 20 °C, and then dissolving them in 20% bleach for counting of retained eggs. Early-stage eggs were quantitated as described (Koelle and Horvitz, 1996). Briefly, age-matched adult animals (24 h past L4 stage) were transferred to a new plate and allowed to lay eggs for 30 minutes. Cell stages of at least 100 laid eggs were analyzed under a Leica M165FC stereomicroscope. Embryos with 1 to 8 cell stages were categorized as “early-stage”, and embryos with >8 cells stages were categorized as “normal stage” eggs.

For acute heat shock experiments, late-L4 animals from strains N2, MIA326, MIA325, and MIA508 were picked onto NGM plates and incubated at 20 °C for 18 hours. Half of the plates were sealed with parafilm and shifted to a 33 °C for 30 minutes, 20 °C for 30 minutes, again shifted back to 33 °C for 30 minutes, and then finally returned to 20 °C for 12 h prior to behavior experiments.

### Microscopy

#### Ratiometric Ca^2+^ imaging

Ca^2+^ activity was performed in freely behaving adult animals at 24-30 h past the late L4 larval stage, as described previously (Ravi et al., 2018a). Worms were mounted between a chunk of NGM and a glass coverslip. Vulval muscle Ca^2+^ activity was recorded through a 20X Apochromatic objective (0.8 NA) mounted on an inverted Zeiss Axio Observer.Z1. A Zeiss Colibri.2 LED illumination system was used to excite GCaMP5 at 470 nm and mCherry at 590 nm for 10 msec every 50 msec. GFP and mCherry fluorescence emission channels were separated using a Hamamatsu W-VIEW Gemini image splitter and recorded simultaneously for 10 min with an ORCA-Flash 4.0 V2 sCMOS camera at 256 / 256-pixel resolution (4x4 binning) at 16-bit depth. The motorized stage was manually controlled using a joystick to maintain the freely behaving animal in the field of view. HSN Ca^2+^ activity was recorded through a 20x Apochromat objective (0.7 NA) on an inverted Leica TCS SP5 confocal microscope using the 8 kHz resonant scanner at 20-30 fps at 256 by 256-pixel resolution, 12-bit depth at 2-4X digital zoom. GCaMP5 and mCherry fluorescence were excited using the 488 nm and 561 nm laser lines, respectively. Animals were recorded until each entered an egg-laying active state, defined as the period one minute before the first egg-laying event and ending one minute after the last egg-laying event. The 10-minute (12,000 frame), two-channel image sequences, centered on the first egg-laying event observed were recorded for subsequent ratiometric analysis. In recordings of animals expressing GFP-C3T in the HSNs, sequences were centered on the first egg-laying event observed within the 15-minute recording period. If no eggs were laid in 15 minutes, the first 10 minutes of the recording were saved. Image sequences were exported to Volocity software (Quorum Technologies Inc.) for segmentation and ratiometric analysis. Ca^2+^ transient peaks from ratio traces were detected using a custom MATLAB script, as described (Ravi et al., 2018a).

To obtain Z-stack images of the HSN neurons in animals expressing GCaMP5 and mCherry alone or along with GFP-C3T, 30 h post-L4 animals were immobilized with 3 μL of 100 mM sodium azide on a 1% agarose pad on a microscope slide. The worms were then covered with a glass coverslip and transferred to the stage of an inverted Zeiss LSM 880 microscope. Z-stack images were recorded with 512x512 pixel resolution, 16-bit depth, and 1.5X digital zoom through a plan-apochromat 63x/1.4 oil objective. GFP and mCherry fluorescence were excited using a 488 nm and 594 nm laser lines, respectively. Image slices were then exported to Volocity software (Quorum Technologies Inc.) for 3D visualization. For HSN morphology analysis, twelve Z-stacks containing a clearly visible HSN cell body, distinguishable presynaptic termini located on either side of the vulval slit, and clear, connecting processes in between cell body and termini were characterized as normal. All others were deemed abnormal.

### Experimental design and statistical analysis

Sample sizes for behavioral assays followed previous (Chase and Koelle, 2004; Collins et al., 2016). Statistical analysis was performed using Prism v.8 or v.9 (GraphPad). Ca^2+^ transient peak amplitudes, widths, and inter-transient intervals were pooled from multiple animals (typically ≥ 10 animals per genotype). All statistical tests were corrected for multiple comparisons (Bonferroni for one-way ANOVA or Fisher’s exact tests; Dunn’s correction for Kruskal–Wallis tests). Each figure legend indicates individual p-values with p<0.05 being considered significant except where indicated otherwise. For the analysis of HSN morphology in transgenic animals expressing mCherry +/- GFP-C3T, mean proportions per genotype of animals with “normal” HSN morphology were compared by Fisher’s exact test. 95% confidence intervals for all mean proportions indicated were calculated using the Wilson score method without continuity correction (Newcombe, 1998).

## Results

### Rho GTPase promotes egg-laying behavior downstream of Trio RhoGEF and independent of circuit development

We have previously shown that Trio RhoGEF acts in both neurons and muscle cells to regulate egg-laying behavior, likely through the PIP_2_/DAG pathway (Dhakal et al., 2022). Previous studies have found that manipulation of Rho signaling using constitutively active Rho-1(G14V) under heat shock promoter or cholinergic neuron-specific promoter increases egg-laying behavior (McMullan and Nurrish, 2011), suggesting Rho-1 regulates synaptic transmission (**Figure 1A**). To test if Rho acts downstream of the Gα_q_-Trio pathway, we transgenically expressed GTP-locked Rho-1 mutant (G14V) to increase Rho-1 activity in wild type or *unc-73(ce362)* Trio loss-of-function mutant animals. Because Rho signaling is tightly regulated and alterations that cause either gain- or loss-of RHO-1 signaling cases lethality (Jantsch-Plunger et al., 2000; McMullan and Nurrish, 2011), we expressed Rho-1(G14V) from a heat shock promoter to increase Rho signaling acutely after animals had reached adulthood (**Figure 1B**) when circuit development was already complete (Ravi et al., 2018b).

**Figure 1.**
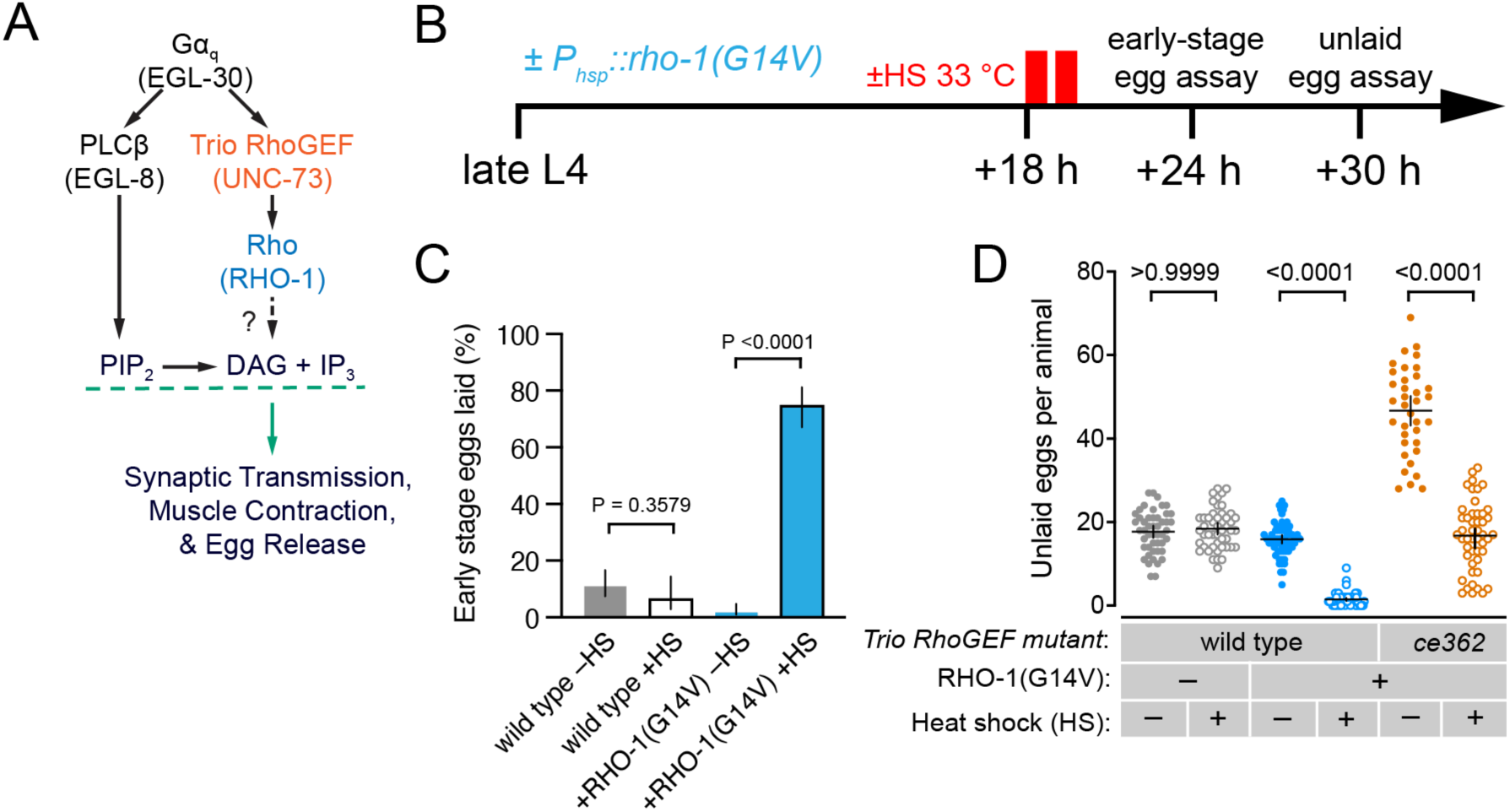
Rho GTPase regulates egg-laying behavior independently of egg-laying circuit development. **(A)** Cell signaling pathways in neurons and muscles that regulate egg laying. **(B)** Experimental procedure for conditional overexpression of Rho-1(G14V) from a heat shock promoter. Late-L4 animals were incubated for 18 hours at 20 °C, half were shifted twice to 33 °C for 30 minutes, and back to 20 °C. Changes in egg laying were then measured by quantifying the stages of eggs laid and then by measuring the number of embryos retained in the uterus. **(C)** Bar plots showing the percentage of early-stage (premature eggs) laid by wild-type and transgenic animals expressing nothing or Rho-1(G14V) from the heat shock promoter ± heat shock induction. Bar indicates percent of early-stage eggs ±95% confidence intervals. Significant p-values are shown; n.s.= not significant (p>0.05, Fisher’s exact test with Bonferroni correction for multiple comparisons; n≥100 eggs per genotype and condition). **(D)** Scatterplot of egg accumulation in wild-type and *unc-73(ce362)* mutant transgenic animals expressing nothing or Rho-1(G14V) ± heat shock induction. The horizontal line indicates mean accumulated eggs ±95% confidence intervals. p-values are indicated (one-way ANOVA with Bonferroni’s correction for multiple comparisons, n≥30 animals per genotype and condition).

After providing heat shock, we first measured the stage of eggs laid as a measure of steady-state egg laying. Wild-type animals typically retain fertilized embryos in the uterus for sufficient time that they reach the 50 to 100-cell stage, whereas animals with hyperactive egg-laying typically lay embryos more quickly, having reached a developmental stage of eight cells or fewer (Chase and Koelle, 2004; Ségalat et al., 1995). Heat shock expression of the constitutively active Rho-1(G14V) mutant strongly and significantly stimulated egg-laying behavior in wild-type animals (**Figure 1C**), with animals releasing more than 70% early-stage eggs while wild-type control animals laid only 10-15% at early stages, like that seen in animals without heat shock. Second, we measured the steady-state accumulation of embryos in the uterus as a proxy for changes in egg-laying behavior. Animals acutely expressing Rho-1(G14V) accumulated significantly fewer eggs (1.5±0.4) compared to heat-shocked control animals which retain ∼17±1 embryos, like that seen in animals without heat shock (**Figure 1D**). Animals bearing a missense mutation in the RhoGEF domain of Trio, *unc-73(ce362)* show strong egg-laying behavior defects (Dhakal et al., 2022; Williams et al., 2007), accumulating more than 45 eggs in the uterus (**Figure 1D**). Heat shock expression of Rho-1(G14V) rescued egg-laying behavior to Trio RhoGEF mutants, resulting in near wild-type retention of ∼17±2 eggs (**Figure 1D**). Together, these results confirm that Rho GTPase promotes egg-laying behavior in *C. elegans,* and it does so downstream of Trio RhoGEF.

### Rho GTPase acts in the HSNs and vulval muscles to promote egg-laying behavior

Our previous experiments confirm Rho-1 promotes egg laying, but whether and where Rho-1 acts within the egg-laying circuit was not clear. To address the site of Rho-1 action, we used cell-specific promoters to dominantly activate or interfere with Rho-1 signaling in the HSNs, the serotonergic command neurons that stimulate contractility of the vulval muscles in the egg-laying circuit (**Figure 2A**). We found that transgenic expression of Rho-1(G14V) in HSNs of otherwise wild-type animals activates egg laying (**Figure 2B**), with 70% of their eggs being laid at early-stages (**Figure 2C**) and a corresponding decrease in egg retention (∼3±1 eggs) (**Figure 2D)**. Conversely, animals expressing a putative dominant-negative Rho-1(T19N) mutant in HSN predicted to be in the GDP-locked form (Santos et al., 1997) accumulated an average of 27±2 eggs, a significant increase (**Figure 2D**). Together, these results confirm that Rho-1 functions in the HSNs to regulate egg-laying behavior in *C. elegans*.

**Figure 2:**
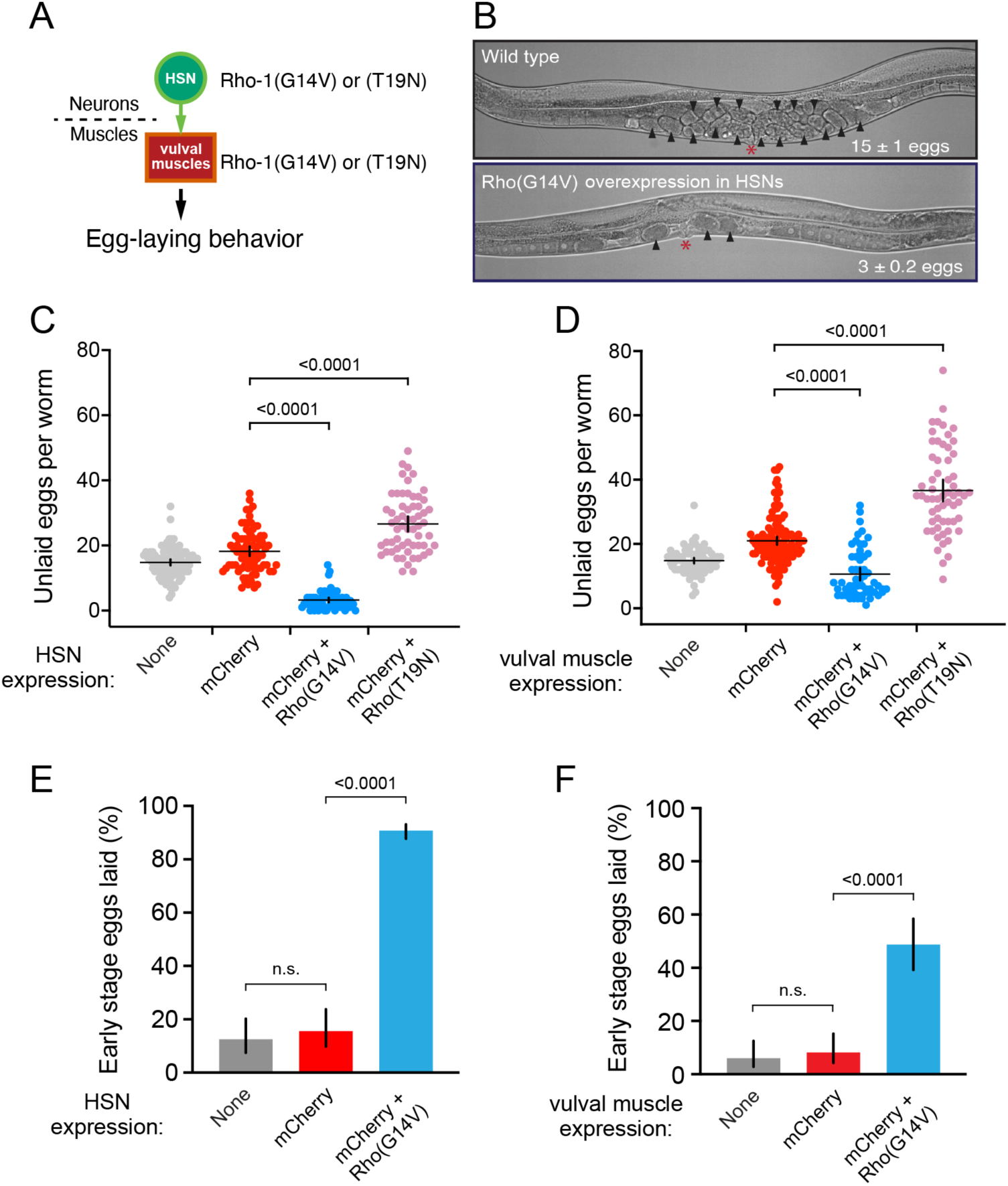
Rho GTPase acts in the HSNs and vulval muscles to promote egg-laying behavior. **(A)** Cartoon of experiment. Rho-1(G14V) or Rho-1(T19N) is expressed in HSN or vulval muscle and egg-laying behavior is measured. (**B**) Micrographs of wild-type animals showing normal egg accumulation (top) and transgenic animals expressing Rho-1(G14V) in HSN showing decreased egg accumulation. **(C)** Scatterplot of egg accumulation in wild-type animals expressing nothing or mCherry with or without Rho-1(G14V) or Rho-1(T19N) in HSN. The horizontal line indicates mean accumulated eggs ±95% confidence intervals. **(D)** Scatterplot of egg accumulation in wild-type animals expressing nothing or mCherry with or without Rho-1(G14V) or Rho-1(T19N) in vulval muscle. The horizontal line indicates mean accumulated eggs ±95% confidence intervals. **(E)** Bar plots showing the percentage of early-stage eggs laid by wild-type animals expressing nothing (grey), mCherry alone (red), or mCherry along with Rho-1(G14V) in HSN (blue). Vertical lines indicate 95% confidence intervals of the mean proportions. **(F)** Bar plots showing the percentage of early-stage eggs laid by wild-type and transgenic animals expressing nothing (grey), mCherry alone (red), or mCherry plus Rho-1(G14V) (blue) in vulval muscle. Vertical lines indicate 95% confidence intervals of the mean proportions. p-values indicated were determined by one-way ANOVA (C-D) or Fisher’s exact test (E-F) with Bonferroni’s correction for multiple comparisons. n.s. indicates p-value >0.05. n≥30 animals per genotype (C-D); n>100 eggs per genotype (E-F).

To examine Rho-1 function postsynaptically, we used a vulval muscle-specific promoter to express Rho-1(G14V) and measured egg accumulation in these animals. We observed that transgenic expression of Rho-1(G14V) in otherwise wild-type animals significantly increased egg laying (**Figure 2E**) leading to 48% of the eggs being laid at early stages with a corresponding decrease in egg retention (∼10±2 eggs) (**Figure 2F**). Conversely, animals expressing the Rho-1(T19N) dominant-negative mutant in vulval muscles accumulated an average of ∼37±3 eggs, a significant increase (**Figure 2F**). Together, these results show that Rho-1 also functions in the vulval muscles to regulate egg-laying behavior in *C. elegans*. Transgenic expression of Rho-1(G14V) in the egg-laying vulval muscles showed a weaker effect than expression in the HSNs. Because the Rho-1 transgenes used in these experiments were analyzed as extrachromosomal arrays, with Rho-1 transgenes mosaically, these results support a more general interpretation that Rho-1 functions in both the HSNs and vulval muscles to promote egg-laying. We confirm these results below using integrated transgenes with more stable patterns of expression.

### Rho signals in HSN to promote Ca^2+^ activity

We hypothesized the changes in egg-laying behavior in the animals that express Rho-1 gain- and loss-of-function mutants in HSNs is due to changes in HSN excitability and/or neurotransmitter release. To test this, we performed ratiometric Ca^2+^ imaging in behaving animals expressing Rho-1(G14V) or Rho-1(T19N) in HSN during egg-laying active states. As we previously reported, wild-type HSNs displayed Ca^2+^ transients in bursts during the egg-laying active state with only intermittent Ca^2+^ transient activity during the inactive state (Collins et al., 2016; Ravi et al., 2021). The normal burst pattern of Ca^2+^ activity was lost in animals expressing Rho-1(G14V) in the HSNs converting instead to a more tonic firing pattern with Ca^2+^ transients observed both in and outside of egg-laying active states (**Figure 3A**). This increased Ca^2+^ transient activity seen in Rho-1(G14V)-expressing animals was reminiscent of the increased bursting HSN Ca^2+^ transient activity found in animals lacking inhibitory Gα_o_ signaling (Ravi et al., 2021). Expression of Rho-1(T19N) putative dominant-negative mutant did not strongly affect the pattern of active/inactive state Ca^2+^ transient activity, despite causing a modest inhibition of egg-laying behavior (**Figure 2C**).

**Figure 3:**
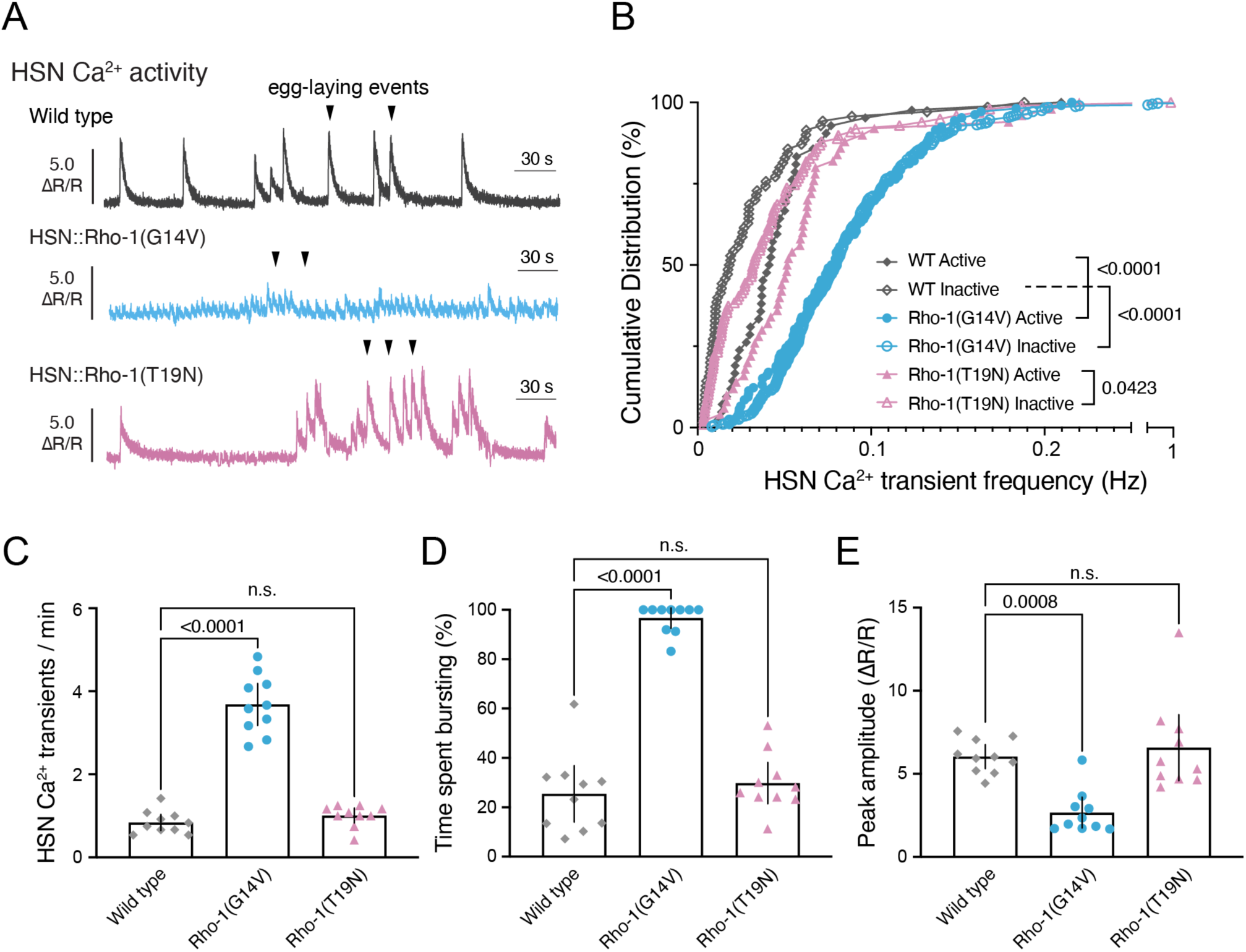
Rho signaling promotes HSN Ca^2+^ activity. **(A)** Representative GCaMP5: mCherry (ΔR/R) ratio traces showing HSN Ca^2+^ activity in freely behaving wild-type (dark grey) or transgenic animals expressing constitutively active Rho-1(G14V) (light blue) or dominant negative Rho-1(T19N) (lavender) mutants in the HSN neurons during an egg-laying active state. Arrowheads indicate egg-laying events; scale indicates 30 s. **(B)** Cumulative distribution plots of instantaneous Ca^2+^ transient peak frequencies in wild-type animals (dark grey diamonds) or transgenic animals expressing constitutively active Rho-1(G14V) (light blue circles), or dominant negative Rho-1(T19N) (lavender triangles) in the HSN neurons. Filled shapes represent instantaneous transient frequencies in the egg-laying active state, which is defined as 60 seconds before and 60 seconds after an egg-laying event, where overlapping egg-laying events are pooled and treated as one. Open versions of each shape represent transients occurring during the egg-laying inactive state. Significant P values are shown; n.s. indicates P>0.05 (Kruskal-Wallis test with Dunn’s correction for multiple comparisons). (**C)** Scatterplots showing average Ca^2+^ transient frequency (per min). Error bars indicate 95% confidence intervals for the mean. Significant p-values are shown; n.s. indicates p >0.05 (One-way ANOVA with Bonferroni correction for multiple comparisons). **(D)** Scatterplots showing average % of time spent bursting or showing high-frequency trains of Ca^2+^ peaks. Significant p-values are shown; n.s. indicates p >0.05 (Kruskall-Wallis test with Dunn’s correction for multiple comparisons). **(E)** Scatterplots showing average Ca^2+^ transient amplitudes (ΔR/R). Significant p-values are shown; n.s. indicates p >0.05 (One-way ANOVA with Šídák’s multiple comparisons test).

To quantify changes in HSN Ca^2+^ activity, we compared instantaneous (**Figure 3B**) and overall (**Figure 3C**) transient frequencies between wild-type and Rho-1 mutants. Animals expressing Rho-1(G14V) had significantly higher Ca^2+^ transient frequencies compared to wild-type or Rho-1(T19N) animals, whether animals were in the egg-laying active state or not (**Figure 3B**). While wild-type animals typically show <1 Ca^2+^ transient per minute, on average, this was increased to more than 5 transients per minute in animals expressing Rho-1(G14V) in the HSNs, a significant difference (**Figure 3C**). Animals expressing Rho-1(T19N) in the HSNs had no significant change in Ca^2+^ transient frequency compared to wild type (**Figure 3C**). We quantified the percent of time animals spent with high frequency Ca^2+^ transients typically seen during the egg-laying active state. Wild-type and Rho-1(T19N) mutant animals typically spend 25-30% of their time in recordings with Ca^2+^ activity seen in bursts (**Figure 3D**), consistent with previous results (Ravi et al., 2021). Animals expressing Rho-1(G14V) in the HSNs spent nearly ∼100% of their time in recordings showing high frequency Ca^2+^ transients typically only seen in egg-laying active states, consistent with the increased egg-laying behavior observed. HSN Ca^2+^ transients in Rho-1(G14V) HSNs were also smaller in amplitude (**Figure 3E**). Whether this reduced amplitude was a consequence of the increased transient frequency or by other changes in HSN excitability is not clear, but a similar activity phenotype was observed in HSNs of animals bearing a gain-of-function mutation the T-type Cav3.1 Ca^2+^ channel homolog, CCA-1 (Zang et al., 2017). Taken together, these results show that increased Rho signaling in HSN potentiates cell excitability, likely driving an elevation of synaptic transmission that promotes vulval muscle excitability and hyperactive egg-laying behavior.

### Rho signaling in HSN promotes vulval muscle Ca^2+^ activity

HSN innervates and provides synaptic input to the vulval muscles (Desai et al., 1988; Li et al., 2013; Ravi et al., 2018b; White et al., 1986). We next tested whether Rho-1(G14V) expression in HSN had consequences on vulval muscle Ca^2+^ activity (**Figure 4A**). As shown in **Figure 4B**, wild-type animals show a two-state pattern of rhythmic vulval muscle Ca^2+^ activity, with infrequent Ca^2+^ ‘twitch’ transients during the inactive state and more frequent twitching and egg-laying Ca^2+^ transients during egg-laying active state (Collins and Koelle, 2013; Collins et al., 2016). In contrast, animals expressing Rho-1(G14V) in HSNs show continuous rhythmic twitching Ca^2+^ transients during both the inactive and active behavior states (**Figure 4B)**. This increased muscle activity was again reminiscent of animals with too little inhibitory Gα_o_ signaling (Ravi et al., 2021) or too much Gα_q_ signaling (Dhakal et al., 2022), consistent with Rho-1(G14V) promoting HSN excitability and neurotransmitter release onto the vulval muscles.

**Figure 4:**
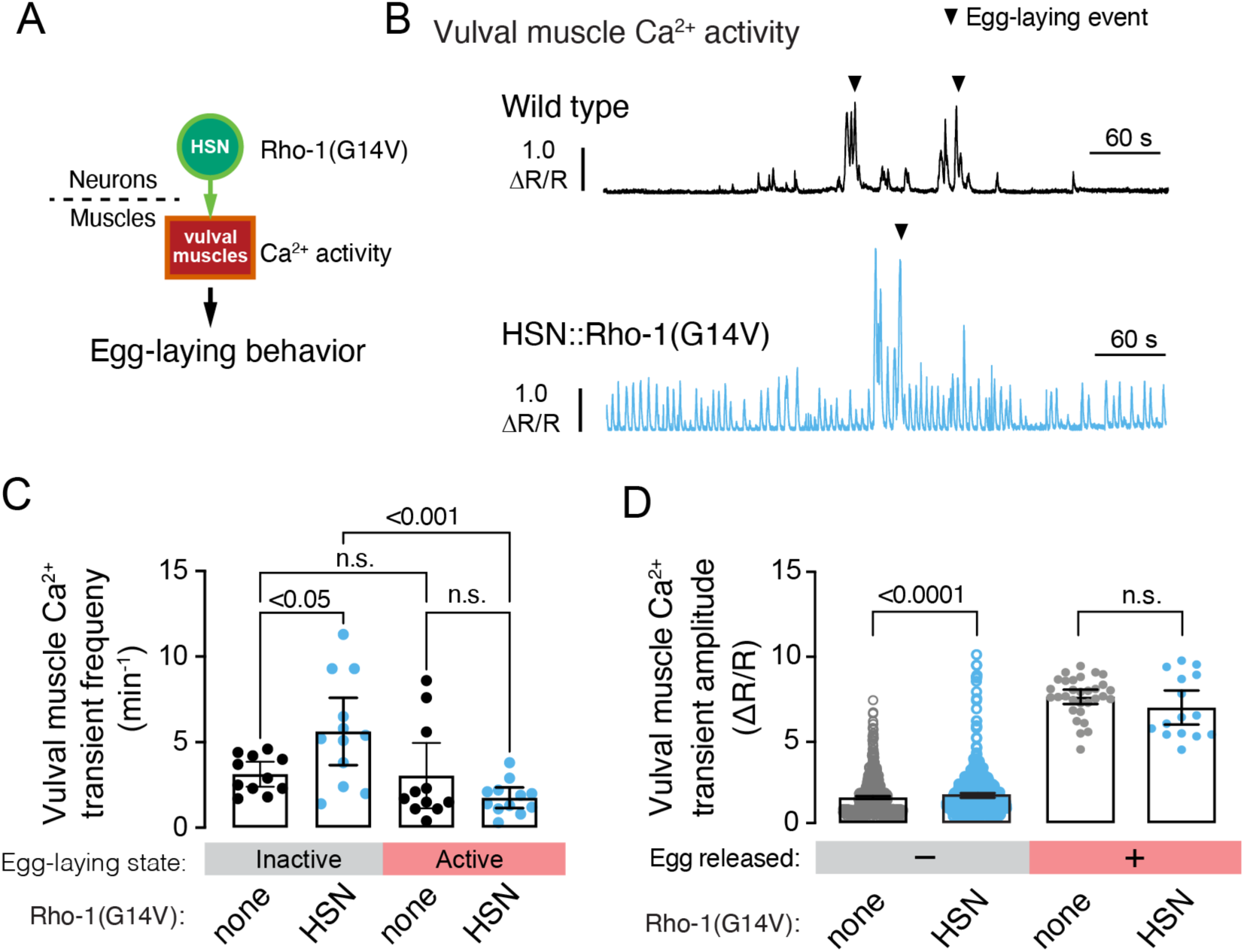
Rho signaling in the HSNs promotes vulval muscle Ca^2+^ activity. **(A)** Cartoon of experiment. Rho-1(G14V) is expressed in HSN and vulval muscle Ca^2+^ activity is measured using GCaMP5. **(B)** Representative GCaMP5:mCherry (ΔR/R) ratio traces showing vulval muscle Ca^2+^ activity respectively in wild-type or in animals expressing Rho-1(G14V) in HSN. Arrowheads indicate egg-laying events. Vertical and horizontal scale bars show GCaMP5/mCherry fluorescence ratio (ΔR/R) and time, respectively. **(C)** Scatterplots showing vulval muscle Ca^2+^ transient frequency in the egg-laying active and inactive states. Line indicates mean eggs laid ±95% confidence intervals; n.s. indicates not significant (p-values are also indicated (one-way ANOVA with Bonferroni’s correction for multiple comparisons; n>10 animals recorded per genotype). **(D)** Scatterplots showing comparisons of vulval muscle Ca^2+^ transient amplitudes during egg-laying or twitch Ca^2+^ transients. The line indicates mean eggs laid ±95% confidence intervals; n.s. indicates not significant (p>0.05; the associated p-value is also indicated; one-way ANOVA with Bonferroni’s correction for multiple comparisons; n>10 animals recorded per genotype).

To further quantify these activity defects, we compared vulval muscle Ca^2+^ transient frequencies and amplitudes in wild-type and Rho-1(G14V)-expressing animals. Expression of Rho-1(G14V) in HSN did not significantly affect vulval muscle Ca^2+^ transient frequency during egg-laying active states, but we did see a significant increase during the inactive state (5.6 ± 2.0 transients per minute) compared to wild-type animals (3.0 ± 1.9 transients per minute) (**Figure 4C**). The increase in vulval muscle activity outside the active state likely results from tonic HSN Ca^2+^ activity in Rho-1(G14V)-expresing HSNs (**Figure 3A**), promoting release of 5HT and NLP-3 neuropeptides which increases vulval muscle excitability (Brewer et al., 2019; Butt et al., 2024; Olson et al., 2023). The increased vulval muscle Ca^2+^ activity may also reflect increased muscle excitability that potentiates rhythmic ACh excitation in phase with locomotor body bends (Collins and Koelle, 2013; Collins et al., 2016; Kopchock et al., 2021). Consistent with this, while egg-laying vulval muscle Ca^2+^ transient amplitudes were not significantly different in animals expressing Rho-1(G14V) in the HSNs (7.6 ΔR/R, wild type vs. 6.9 ΔR/R, G14V), the smaller rhythmic twitching Ca^2+^ transients were significantly larger (1.7 ± 0.1 ΔR/R) compared to wild type (1.5 ± 0.1 ΔR/R, **Figure 4D**). Together, these results show that Rho signaling in HSN potentiates vulval muscle activity, consistent with Rho-1 acting presynaptically to promote neurotransmitter release.

Because transgenic expression of the Rho-1(T19N) mutant only partially inhibited egg laying and had no effect on HSN Ca^2+^ activity, we reasoned this may be caused by insufficient inhibition of Rho-1 function. To address this, we expressed C3 Transferase (C3T) to more completely inactivate Rho-1 signaling (Chardin et al., 1989; McMullan et al., 2006). Transgenic expression of C3T in the HSNs from the *tph-1* promoter (**Figure 5A**) caused a strong and significant increase in egg accumulation (**Figure 5B-C)**, reflecting a block in egg laying. Animals bearing either of two independently integrated transgenes accumulated ∼45 eggs in the uterus (**Figure 5D**), similar to animals bearing the *egl-6(n582gf)* mutation with too much inhibitory Gα_o_ signaling in HSNs (Ringstad and Horvitz, 2008). C3T expression caused significantly more egg accumulation than animals expressing Rho-1(T19N) (see **Figure 2C**). Because Rho-1 regulates key developmental processes, it was possible that complete Rho-1 inhibition by C3T could impair HSN development. To address this, we used confocal microscopy to visualize HSN morphology in animals expressing C3T. We found that HSN cell body placement and neurite morphology was grossly normal and similar to non-transgenic control animals (**Figure 5E-G**), consistent with the onset of HSN expression from the *tph-1* promoter occurring after HSN development is complete and egg-laying events have begun (Ravi et al., 2018b; Tanis, 2008; Tanis et al., 2008). Together, these results indicate Rho inhibition in HSN strongly blocks egg-laying behavior, and that this inhibition is not caused by gross alterations in HSN development.

**Figure 5:**
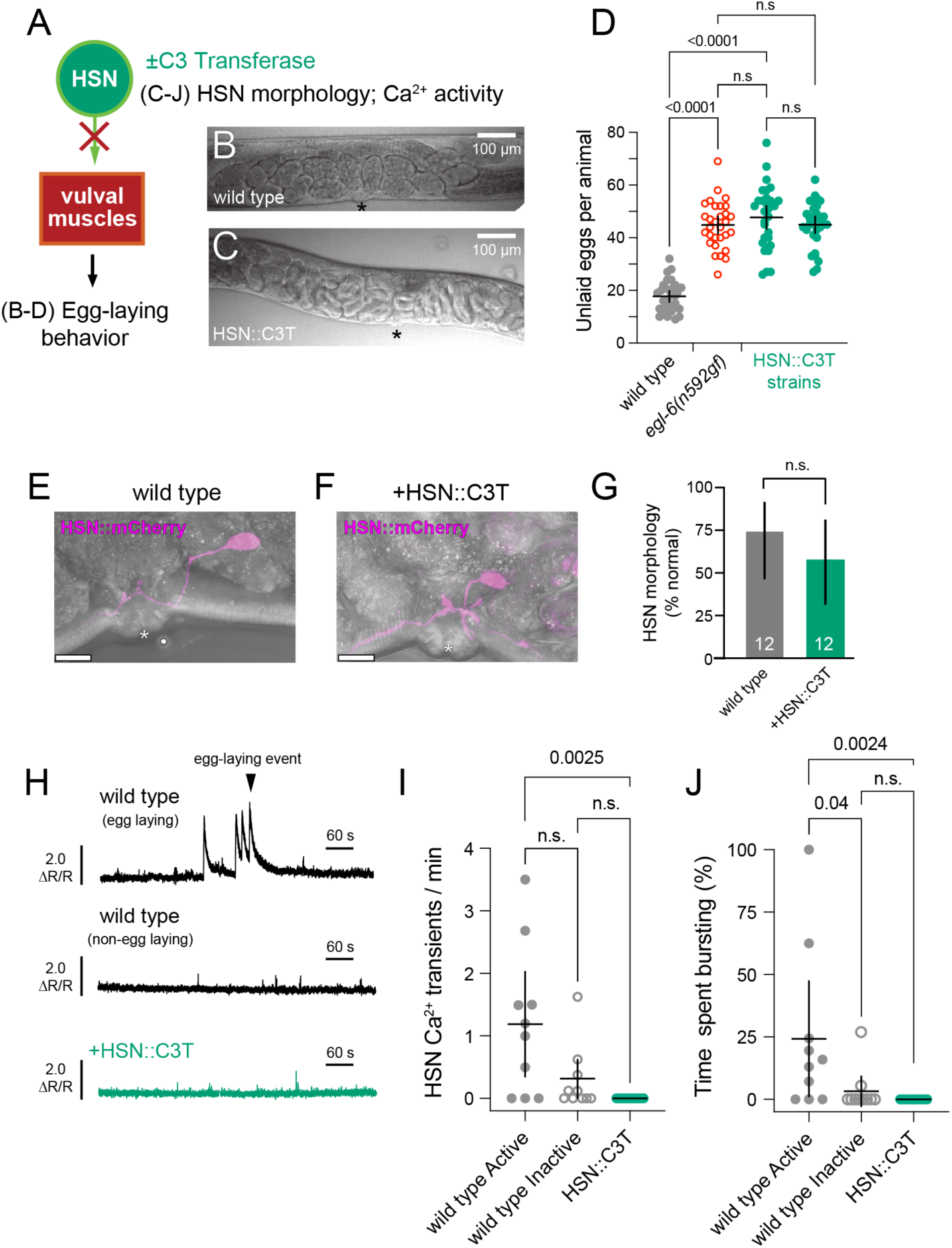
Rho signaling in the HSNs is required for HSN Ca^2+^ activity and normal egg-laying behavior. (**A**) Cartoon of the experiment. C3 transferase (C3T) is expressed in the HSNs and changes in egg-laying behavior are measured along with HSN Ca^2+^ activity. (**B**) Micrograph of wild-type animal showing normal egg accumulation. (**C**) Micrograph of transgenic animal expressing C3T in the HSNs showing increased egg accumulation. (**D**) Scatterplot of unlaid eggs accumulated in the uterus in wild-type (grey), *egl-6(n592gf)* mutants (red open circles), and two independent transgene integrants expressing C3T and mCherry in the HSNs from the *tph-1* promoter (green). (**E-F**) Representative fluorescence micrographs of animals expressing mCherry and/or GFP-C3T in the HSN neurons. Asterisks indicate the approximate location of the vulval opening. Scale bars are 10 μm. (**G**) Bar graph showing proportions of wild-type and C3 transferase-expressing animals with grossly normal HSN morphology. Error bars show 95% confidence intervals for the mean proportion. P>0.05, Fisher’s exact tests. (**H**) Representative HSN Ca^2+^ activity traces in wild-type animals with and without egg-laying events or transgenic animals expressing C3T in the HSNs. **(I)** Scatterplot of frequencies of HSN Ca^2+^ transients of wild-type animals in active and inactive egg-laying behavior states or in C3T-expressing mutants. (**J**) Scatterplot of time spent by recorded animals with HSNs showing bursting activity (peak intervals ≤30 s) in wild-type active (grey filled) or inactive (grey open circles) egg-laying behavior states of wild-type animals, or in C3T-expressing mutant animals (green filled circles). P-values displayed above comparison brackets were determined by (B) one-way ANOVA with Bonferroni multiple comparisons test or (F-G) Kruskall-Wallis test with Dunn’s correction for multiple comparisons; n.s. indicates not significant (P>0.05). Number of animals: (D) n = 30; (E-G) n=12; (H-J) n = 10.

To determine how Rho-1 inhibition blocks HSN neurotransmitter release, we performed ratiometric Ca^2+^ imaging in HSNs of C3T-expressing animals. We found HSN Ca^2+^ transient activity was largely eliminated upon C3T expression, resembling Ca^2+^ activity seen in wild-type animals during the inactive behavior state (**Figure 5H**). Wild-type animals during the active state typically have ∼1 HSN Ca^2+^ transient per minute, most of which come in ‘bursts’ during egg-laying active states. We observed zero Ca^2+^ transients in C3T-expressing animals (**Figure 5I**), consistent with HSN excitability being dramatically inhibited. Measurement of ‘bursting’ activity showed a similar effect, with C3T-expressing animals spending 0% of time in the bursting state compared to the 25% of time spent by wild-type animals (**Figure 5J**). This inhibition of HSN Ca^2+^ activity upon Rho-1 inhibition is strikingly similar to (although stronger than) the effects seen upon expression of the GTP-locked Gα_o_ or in *egl-10* mutant animals lacking the Gα_o_-directed RGS protein that terminates its signaling (Ravi et al., 2021). These results show that Rho-1 function is required for HSN Ca^2+^ transient activity, likely because Rho signaling promotes HSN excitability.

HSN promotes vulval muscle activity and contractility for egg laying. We predicted that loss of HSN Ca^2+^ activity in animals expressing C3T would also have reduced vulval muscle Ca^2+^ activity, reflecting a defect in HSN synaptic transmission (**Figure 6A**). While egg-laying events were much less frequent in C3T-expressing animals compared to wild-type controls (data not shown), we still observed significant vulval muscle Ca^2+^ transient activity and egg laying in animals expressing C3T in the HSNs (**Figure 6B**). These results resemble the vulval muscle Ca^2+^ activity phenotypes of animals lacking HSNs altogether (Collins et al., 2016). Measuring vulval muscle Ca^2+^ transient amplitudes, we find egg-laying transients were not significantly different between wild-type and HSN::C3T-expressing animals (∼2.5 ΔR/R; **Figure 6C**). We also observed that C3T-expressing animals showed similar vulval muscle twitch Ca^2+^ transient amplitudes whether they were in the egg-laying active or inactive state (0.71 ±0.1 ΔR/R vs. 0.72 ±0.04 ΔR/R; **Figure 6C**). This contrasts with wild-type animals where twitch transients are significantly stronger during the egg-laying active state (active: 0.85 ±0.06 ΔR/R vs. inactive: 0.62 ±0.07 ΔR/R; **Figure 6C**). This difference largely resulted from a lack of twitch transients around egg-laying events in C3T-expressing animals (**Figure 6B**). As shown in **Figure 6D**, twitch Ca^2+^ transient frequency increased during the egg-laying active state in wild-type animals (median 0.2 s^-1^ vs. 0.1 s^-1^), but this was not seen in C3T-expressing animals (median 0.13 s^-1^ vs. 0.14 s^-1^). As Rho-1 inhibition by C3T completely blocks HSN Ca^2+^ activity (**Figure 5H**), we interpret these results as showing vulval muscle Ca^2+^ activity in C3T expressing animals is largely HSN independent. Instead, egg laying in these animals appears to be driven by stretch-dependent activation of the vulval muscles, consistent with our previous results (Collins et al., 2016; Medrano and Collins, 2023; Ravi et al., 2021). Together, our results show that Rho-1 signaling promotes cell excitability and Ca^2+^ activity. Rho-1 signaling promotes neurotransmitter release which subsequently stimulates postsynaptic cells to drive behavior. Inactivation of Rho-1 function largely eliminates presynaptic cell excitability and neurotransmitter release.

**Figure 6:**
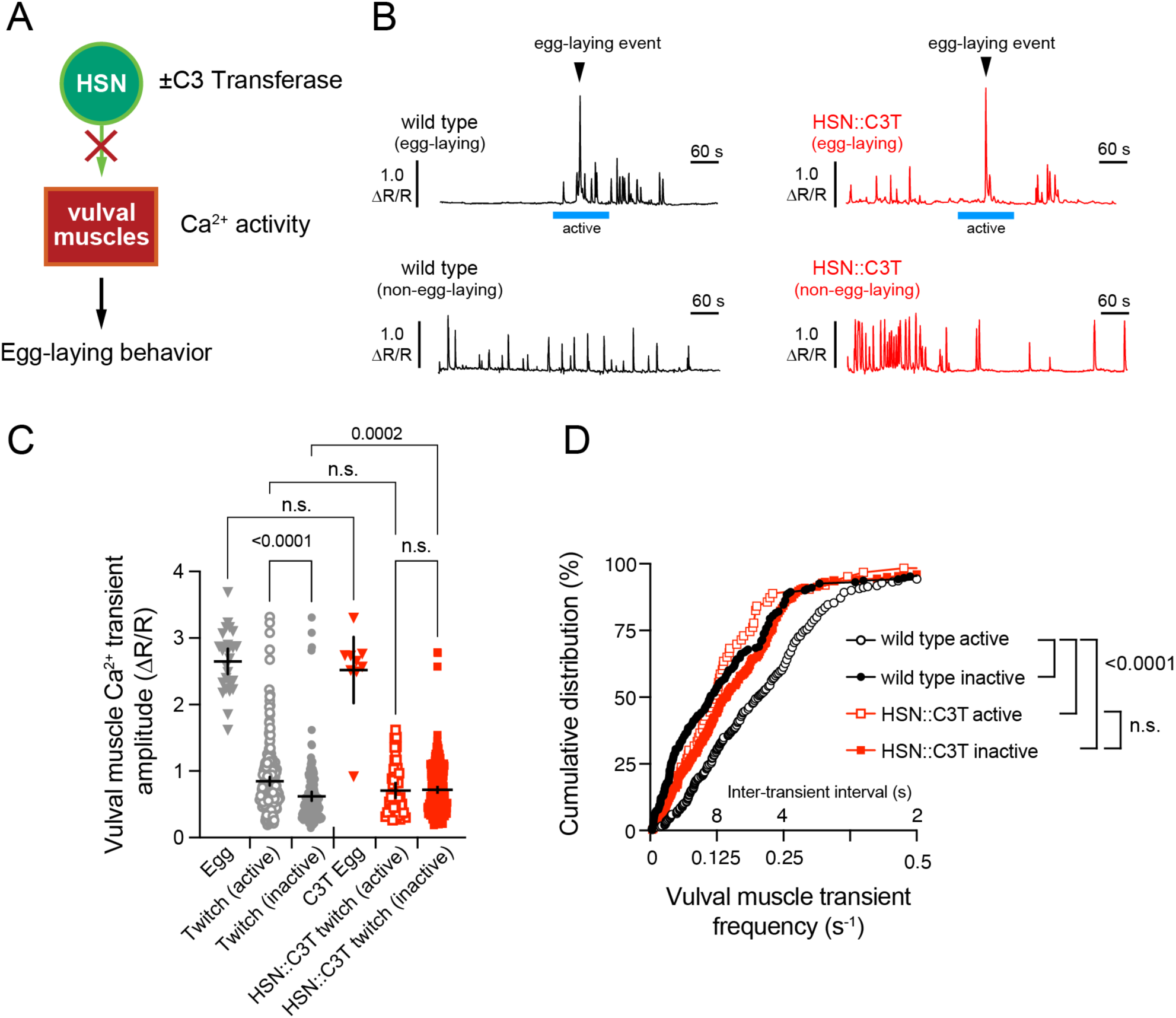
Rho signaling promotes HSN neurotransmission and vulval muscle Ca^2+^ activity around egg-laying behavior states. **(A)** Cartoon of the experiment. C3 transferase (C3T) is expressed in HSN and vulval muscle Ca^2+^ activity is measured using GCaMP5. **(B)** Representative traces of vulval muscle Ca^2+^ activity in wild-type or C3T-expressing animals. Blue bars indicate egg-laying active states operationally defined as one minute before and one minute after egg-laying events (arrowheads). **(C)** Scatterplot of vulval muscle Ca^2+^ peak amplitudes in wild-type (grey) and mutant (green) animals during egg-laying events or vulval muscle twitches observed during the active or inactive egg-laying behavior states. Symbols indicate individual transient amplitudes; error bars indicate the mean ±10% confidence interval. **(D)** Cumulative distribution plot showing instantaneous vulval muscle Ca^2+^ transient frequencies in wild-type (black circles) or C3T-expressing animals (red squares) during the active (open symbols) or inactive (filled symbols). Data collected from 10 animals per genotype. P-values displayed above comparison brackets were determined by Kruskal-Wallis test with Dunn’s test for multiple comparisons.

## Discussion

Using molecular genetics and Ca^2+^ imaging techniques we characterized how Rho GTPase signaling regulates synaptic transmission in the *C. elegans* egg-laying circuit. Our data show that Rho-1 signals downstream of Trio to promote egg laying. Rho-1 functions both presynaptically in the HSNs and postsynaptically in the vulval muscles to promote egg laying, likely acting in part to promote cell excitability. Thus, Rho-1 function is both necessary and sufficient for activation of HSNs and their subsequent excitability of the vulval muscles for proper egg-laying behavior.

Our results show that changes in excitatory Rho-1 signaling have strikingly similar, but opposite, effects as inhibitory Gα_o_ signaling. Expression of Rho-1(G14V) in HSNs drives tonic HSN Ca^2+^ activity, increased vulval muscle Ca^2+^ activity, and stimulates egg laying–phenotypes that resemble the effects when inhibitory Gα_o_ signaling is blocked (Ravi et al., 2021) or excitatory Gα_q_ signaling is increased (Dhakal et al., 2022). Conversely, inactivation of Rho-1 signaling strongly inhibits HSN Ca^2+^ activity and egg laying, phenotypes associated with too much inhibitory Gα_o_ signaling or disrupted Gα_q_ signaling (Koelle and Horvitz, 1996; Ravi et al., 2021; Tanis et al., 2008). We have previously shown that Gα_o_ signaling stabilizes the HSN cell membrane potential (Ravi et al., 2021), and we propose that Gα_q_ signaling through Rho electrically excites cells, antagonizing inhibition by Gα_o_ and its putative effectors. This increase in excitability and Ca^2+^ activity by Gα_q_ and Rho signaling would be expected to enhance the synaptic localization of the vesicle release machinery, facilitating neurotransmitter release from neurons like the HSNs (Lackner et al., 1999; McMullan et al., 2006). Conversely, stabilization of the membrane potential by inhibitory Gα_o_ signaling would be predicted to reduce excitability and subsequent localization of the synaptic vesicle release machinery (Nurrish et al., 1999). Our previous work showing that phorbol esters rescue Ca^2+^ activity and egg-laying defects of Gα_q_ and Trio signaling mutants suggested that DAG production may be a major consequence of Gα_q_ signaling (Dhakal et al., 2022). Our work showing similar Ca^2+^ activity effects by increased Rho-1 signaling further suggests a connection between Rho-1 and DAG signaling.

How would DAG modulate cell excitability and Ca^2+^ activity? One prediction would be that DAG interacts with its prototypical effectors like UNC-13 and Protein Kinase C, containing the C1 domain to stimulate cell Ca^2+^ transient activity (Lackner et al., 1999). UNC-13 is an important regulator of synaptic vesicle fusion that requires calcium influx into the cell (Richmond et al., 1999), but whether PKC is a second target and how it might regulate synaptic excitability is not yet clear. Phorbol esters still stimulate egg laying in animals lacking UNC-13 or PKC orthologs (Dhakal et al., 2022), suggesting DAG acts in part through other targets. DAG can directly modulate the gating of ion channels which might directly promote cell excitability. Several ion channels such as canonical TRP cation channels (TRPC) are activated by DAG (Hofmann et al., 1999). In *C. elegans*, KCNQ family members show sensitivity to DAG-mimetic phorbol esters (Wei et al., 2005). Two-pore domain TASK potassium channels can also be regulated by DAG, suggesting DAG mediates K^+^ channel closure in response to Gα_q_ signaling (Wilke et al., 2014), raising cell excitability. Previous work has shown a role for ion channels like NALCN (NCA-1 and NCA-2) downstream of Gα_q_ and Rho signaling (Topalidou et al., 2017a; Topalidou et al., 2017b). However, we have found that NALCN channels are largely dispensable for Gα_q_ and Gα_o_ modulation of egg laying (Jose and Collins, 2023). T-type Ca^2+^ channels regulate egg laying downstream of G protein signaling (Zang et al., 2017). These channels can be modulated by bioactive lipids like arachidonic acid which are synthesized from DAG precursors (Barbara et al., 2009; Chemin et al., 2014). As T-type Ca^2+^ channels mediate a ‘window current’ whose bistability regulates burst firing (Crunelli et al., 2004), their function and modulation by bioactive lipids downstream of G protein signaling would contribute to the two-state firing behavior of the HSNs (Ravi et al., 2021; Waggoner et al., 1998).

Trio physically interacts with both presynaptic and postsynaptic components where it promotes excitatory transmission. Trio was first identified as a direct interaction with the LAR transmembrane tyrosine phosphatase (Debant et al., 1996). Subsequent work showed Trio’s N-terminal spectrin repeats interact with presynaptic proteins Bassoon and Piccolo (Terry-Lorenzo et al., 2016). Bassoon binds to other presynaptic components where it helps organize presynaptic Ca^2+^ channels (Altrock et al., 2003; Davydova et al., 2014) which might contribute to the changes in cell Ca^2+^ activity we observe. Bassoon and Piccolo also interact with proteins that regulate actin assembly (Fenster et al., 2003; Kim et al., 2003; Wang et al., 1999). We do not see dramatic effects on HSN morphology and presynaptic structure upon Rho inactivation by C3-Transferase, suggesting the effects on behavior are not simply a consequence of synapse elimination. That we see dramatic changes in HSN Ca^2+^ activity upon alterations of Rho-1 signaling are similarly consistent with an direct role for Rho in regulating cell excitability. Trio is also a target of postsynaptic CaMKII phosphorylation where it promotes plasticity (Herring and Nicoll, 2016). Like loss of DGK-1, loss of *C. elegans* CaMKII causes behavior phenotypes that resemble too much Gα_q_ signaling (Reiner et al., 1999), suggesting CaMKII may have conserved targets in *C. elegans*. Trio and Rho also have other known interactors whose functions in the egg-laying circuit remain under-explored, including Rho Activated Kinase (Ishizaki et al., 1996; Leung et al., 1995; Matsui et al., 1996). Together, these results support a model where Gα_q_ activation of Trio and Rho occurs at a larger synaptic protein complex, coordinating changes in excitability and synaptic transmission, some of which may be driven by local changes in protein phosphorylation that may also affect actin assembly.

## Data Availability statement

All the data, reagents, and strains used in this study are available from the corresponding author upon request.

## Acknowledgements

We thank Drs. Laura Bianchi and Michael Koelle along with members of the Collins and Koelle labs for helpful discussions and feedback on the manuscript. Some of the strains used in this study were provided by the *C. elegans* Genetics Center, which is funded by NIH Office of Research Infrastructure Programs (P40 OD010440).

